# Microstructure of Ingestive Behavior in *Mus Musculus*

**DOI:** 10.1101/529198

**Authors:** Giorgio Onnis, Ethel Layco-Bader, Laurence Tecott

## Abstract

We describe a novel quantitative home cage monitoring (HCM) approach for dissecting spontaneous patterns of ingestive and locomotor behaviors into a hierarchically organized series of behavioral facets or endophenotypes. Fine-grained analyses of a large multimodal 16-strain behavioral dataset collected from 169 mice revealed bouts of feeding, drinking and locomotor behaviors occurring within animals’ Active States. We have automated the detection of these bouts and their discrete properties including bout sizes, rates, durations, and intensities. We have developed a hierarchically organized model of behavioral organization enabling analysis of relationships among Active/Inactive State properties and those of feeding, drinking and locomotor bouts. Robust and analogous patterns of interrelationships among these endophenotypes were found for feeding, drinking behaviors, and these differed markedly from those for locomotor behaviors. For feeding and drinking, patterns of reciprocal relationships were observed for pairs of endophenotypes at multiple hierarchical levels. Moreover, endophenotype variability was highest at lowest hierarchical levels progressively diminished at higher levels, so that variability of gross levels of food and water intake were much less than those of their lower level determinants. By contrast, interrelationships among locomotor endophenotypes differed markedly from those of ingestive behavior. Altogether, these findings raise the possibility that behavioral regulation of food and water intake may make an important contribution to the homeostatic maintenance of energy and volume balance.

## INTRODUCTION

Mouse models have been particularly useful for investigating complex homeostatic processes such as energy and volume balance. They have been particularly valuable for illuminating the regulation of molecular/cellular systems, endocrine and neural pathways that underlie these processes^1-3^. Despite these successes, the use of mouse models has yet to similarly refine an understanding of the extent to which mammals regulate their behavior in the service of homeostatic control. In studies of energy balance, investigators often rely on food intake measurements made outside the home cage in novel enclosures to which the animals have not acclimated. In this case, results may be confounded by novelty-induced effects on exploration and anxiety. Short-term assessments of food intake may fail to take into account the profound impact of circadian rhythms on physiology and ingestive behavior. Behavioral assays that focus narrowly on feeding or narrowly on physical activity fail to take into the account the extent to which distinct behavioral determinants of energy balance interact.

Several of these limitations have been addressed by approaches for monitoring spontaneous home cage behaviors, which offer the advantages of minimizing novelty effects, simultaneous assessment of diverse interacting behaviors, and the ability to capture the impact of circadian influences. The typical application of these approaches are hindered however, by analytical methods that apply arbitrary time bins for the detection of behavioral endpoints for which no standardized definitions have been established^4,5^. The development of a unifying conceptual framework for a principled identification of behavioral endophenotypes and the manner in which they are coordinately regulated by the central nervous system could be invaluable for exploring the contribution of behavioral processes to organismal homeostasis.

We have addressed these issues by developing a quantitative home cage monitoring (HCM) approach for dissecting spontaneous behavioral patterns into a hierarchically organized series of subfeatures (endophenotypes)^6^. The classification scheme is based on the concept that lives of small rodents may often be characterized as a series of alternations between Active States (ASs), excursions from a nest or burrow accompanied by diverse active behaviors (including foraging and ingestion), and Inactive States (ISs) at the nest, during which sleep and rest occur^7-9^. We had recently published a report validating the approach in a survey of 16 genetically diverse inbred strains^10^. Using this as a starting point, we describe here the automated assessment of feeding, drinking and locomotor bouts of shorter time scales that occur within ASs. The simultaneous detection and analysis of these features enables “behavioral dissections” that provide unprecedented precision and sensitivity for behavioral phenotyping.

We report here the discovery of a series of robust relationships among endophenotypic measures of ingestive behavior that provide insight into the manner in which behavioral regulation contributes to the biological robustness of homeostatic systems mediating energy and volume balance. Specifically, we propose mechanisms through which experimental influences on ingestive behavior endophenotypes are compensated for (buffered) by others in ways that minimize their effects of gross measures of intake. This is consistent with the widely-held view that centrality of proper energy balance to survival has favored the evolution of robust redundantly-functioning systems.

## RESULTS

### Feeding and drinking bouts

Ingestive behavior events that occur within ASs are expressed in episodic clusters, or bouts (Fig. 1a). As we had previously done for ASs, we have developed and automated approaches for identifying such feeding bouts, drinking bouts, and their properties. We utilized an approach focused on the temporal clustering of behavioral events that has been productively applied in several species. We examined temporal gaps between feeding or drinking events and connected and established a threshold duration value for feeding and drinking interevent intervals (Feeding Bout Threshold (FBT) and Drinking Bout Threshold (DBT), respectively). Consecutive events with interevent intervals less than the threshold were considered to define bouts occurring in the time intervals defined by the onset of the first and offset of last event within each bout. We assessed the robustness of this approach by examining the impact of a wide range of bout thresholds on bout numbers. Bout designation was robust for all strains, as indicated by marked similarities in bout numbers using bout thresholds ranging from x to x min. These values were used for all subsequent ingestive bout analyses. Optimal hyper-parameters for the Strain Survey Dataset were: FBT = 30s, WBT = 30s,

**Figure 1.**
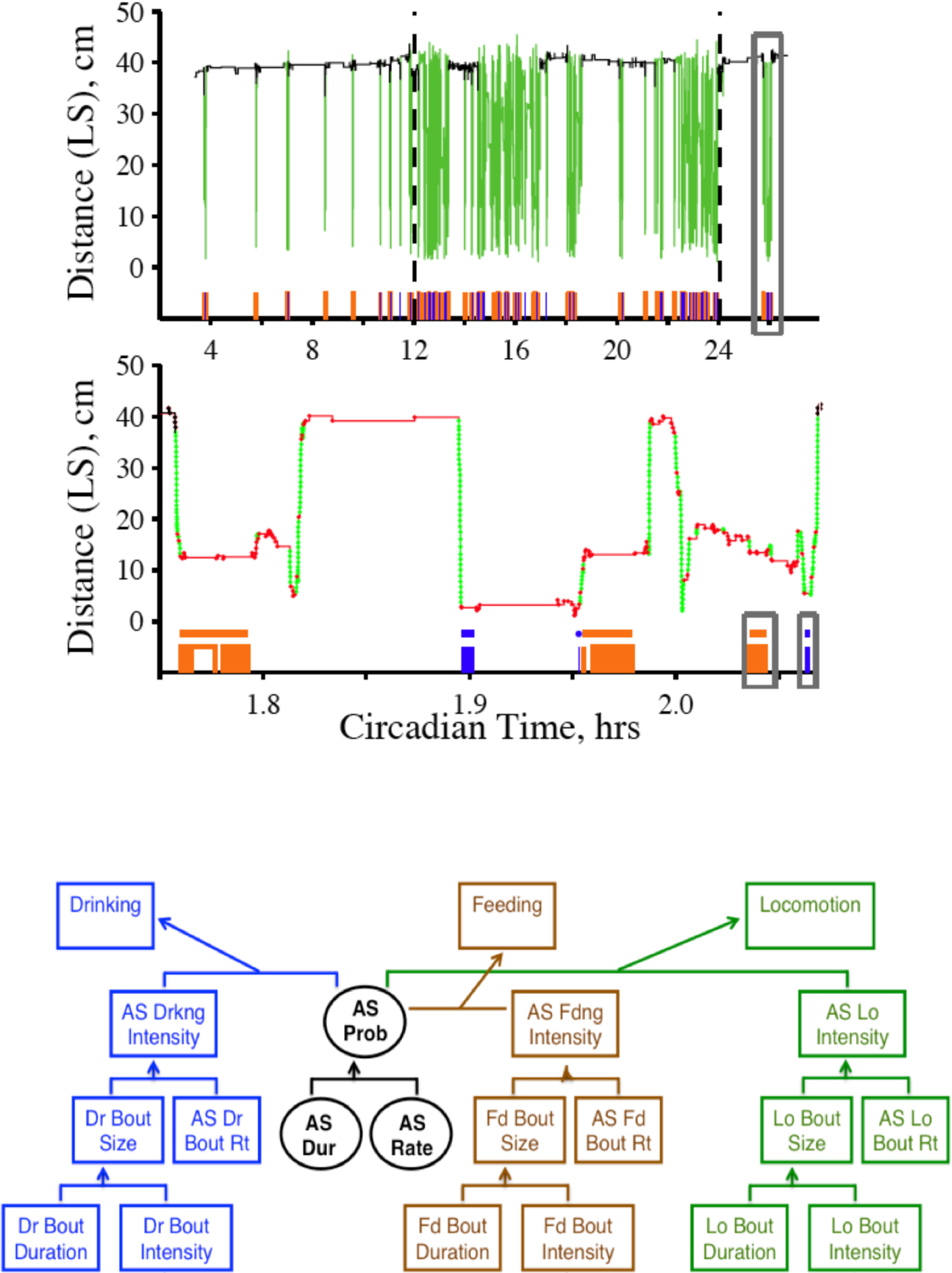
Hierarchical model incorporating active state and bout endophenotypes for feeding, drinking and locomotion. **(a)** The top panel displays a 24 hr behavioral record for a representative C57BL/6J mouse, with locomotor events in green and cage locations expressed on the Y-axis as animals’ distance from the lick spout. Below that, feeding (brown) and licking (blue) events are displayed. Active States in which these behaviors typically co-occur alternate with periods of inactivity at the nest (located 39 cm from lick spout). The last AS in this record is replotted below with an expanded time axis, revealing the output of algorithms that quantify feeding bouts (brown bars) and drinking bouts (blue bars). Locomotor bouts are composed of strings of high-velocity movements, indicated in green. Lower velocity movements are shown in red. **(b)** A framework for describing behavioral organization as a hierarchically organized series of Active State and Bout endophenotypes. The top tier represents gross amounts of food/water intake and locomotor distance. The tier below includes Active State Probability (AS Prob), and Active State Intensities for drinking, feeding, and locomotion. At the next tier are the determinants of Active State Intensities, bout sizes and Active State bout rates. The bottom tier contains determinants of bout sizes, bout durations and intensities.

### Locomotor bouts

Raw spatial and temporal data tracking of the center of mass for each mouse were assessed and movement bout velocity thresholds (MBVTs; movement velocities below which locomotor bouts are considered to have terminated), as well as movement bout distance thresholds (MBDTs; minimum locomotor path distances required to designate movement bouts) were determined as described in Methods. Optimal hyper-parameters for the Strain Survey Dataset were MBVT = 1cm/s, MBDT = 5cm.

### Model of behavioral organization

The ability to parse the behavioral record into ASs and the bouts expressed within them enabled us to develop a hierarchically organized model of behavioral organization incorporating diverse behaviors, each subdivided into behavioral feature components (Fig. 1b) occurring across time scales ranging from days to msec. The top tier represents simple gross measures of ingestion and locomotor distance. These amounts represent the product of ASP and AS intensities for the behavior analyzed, expressed as mass of food/water or distance traveled per unit AS time. In turn, ASPs are the product of AS durations and AS rates (e.g.; number of ASs per day). Likewise, AS intensities are the product of bout sizes and AS bout rates (e.g., rate per hour AS time). Bout sizes are determined by features at the lowest tier, bout durations and intensities (e.g., amount of food consumed or distance traveled per second bout time).

### Reproducibility of HCM data and cohort effects

The utility of this approach for behavioral dissection and its suitability for high-throughput applications is dependent upon the reliability and reproducibility of HCM feature data. It is therefore noteworthy that this strain survey dataset was collected from 7 separate cohorts of mice run over an 11 month period. Individuals from each of the 16 strains were widely distributed across cohorts (most commonly 1-2 animals from each strain were run in each cohort). Since 4 independent cohorts contained a subset of 12 animals with identical strain distributions (C57BL/6J n=1, BALB/cBYJ n=1, A/J n=2, DBA/2J n=2, C3H/HeJ n=1, SWR/J n=2, FVB/NJ n=2 and WSB/Ei n=1), we used their data to assess replicability. Analysis of variance for the 21 model features revealed no effects of cohort as a covariate. Accordingly, very low levels of cohort-to-cohort variability were further indicated by low coefficients of variance for the 21 feature means across these 4 cohorts (average CV: 0.071, Fig. S1).

### Relationships among body weight and levels of feeding, drinking and locomotion

Strain mean body weights ranged from 14.9 +/− 0.4 g (MOLF/Ei, mean +/− standard error) to 37.6 +/− 1.0 g (AKR/J) (Fig. S2). A cluster of strains exhibited body weights less than 20 g, and the remainder of strains were continuously distributed. The rank ordering of strains for body weight and gross levels of ingestion and locomotion were in accord with prior reports^11,12^. Daily mean food intake values ranged continuously from 3.31 +/− 0.10 g (SPRET/Ei) to 5.39 +/− 0.14 g (BALB/cByJ). Daily mean water intake ranged from 2.87 +/− 0.07 g (JF1/Ms) to 5.67 +/− 0.29 g (DBA/2J). Daily mean locomotor distance values were continuously distributed over a wider range (3.6-fold) than ingestive values, ranging from 138 +/− 14 m (C3H/HeJ) to 493 +/− 39 m (WSB). Locomotor data are not presented for 2 of the 16 strains in the study (CZECHII/Ei and CAST/Ei), due to confounding effects of animals climbing and hanging from the undersides of cage lids.

Correlation coefficients for relationships between body weight gross levels of ingestion and locomotor distance were performed using average values for individual mice (Fig. 2). Significant correlations were observed between body weight and food intake (positive, r = 0.685) and locomotor distance (negative, r =- 0.407). No significant correlations were observed between locomotor distance and food or water intake. A well-known positive correlation between food and water intake was (r = 0.653) was also observed. We then determined the extent to which variation in body weights could be accounted for locomotor distance and feeding values. Multiple regression revealed a significant relationship, with food intake and locomotor distance accounting for 80% of the variance on body weight.

**Figure 2.**
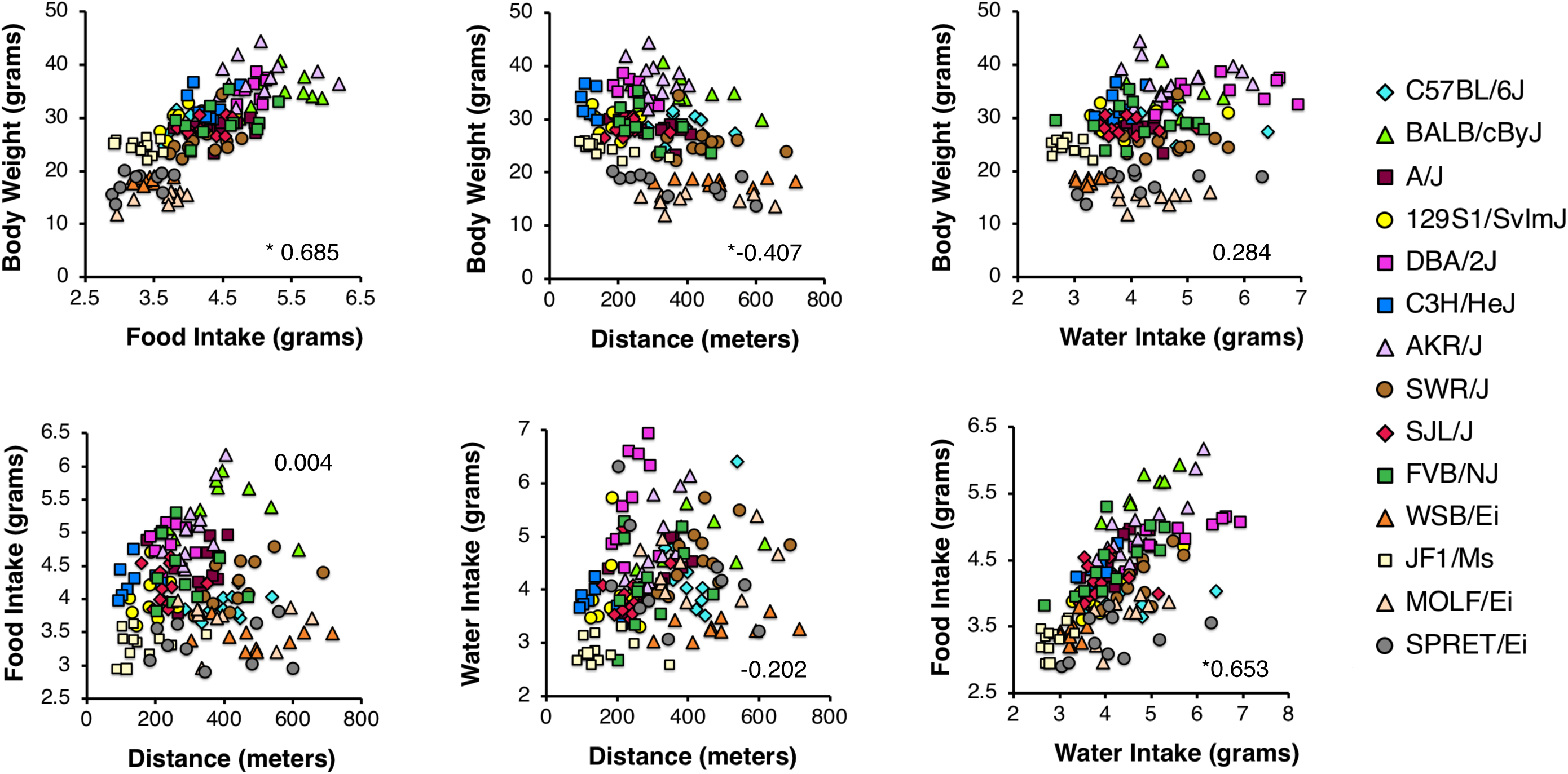
Correlations among body weight, food/water intake and locomotor distance. Scatterplots between feature pairs, with each datapoint representing the 24 hr average feature value for each mouse. Datapoints are color-coded for strain. Pearson correlations coefficient values are shown in each plot. Statistically significant correlations are indicated by asterisks.

### Variability in levels of feeding, drinking and locomotion

Variability in gross daily levels of feeding, drinking and locomotion were assessed at 3 levels: 1) between strains, 2) between individuals within strains, and 3) within individual mice across multiple days. The between-strain coefficient of variance (CV) for locomotor distance was greater than twice that for feeding and drinking (Fig. 3a). We examined within-strain variability by determining CVs for daily average values across individuals of each strain. A consistent pattern generalized across strains: variability in daily food intake was low, with intermediate levels seen for water intake, and highest levels for locomotor distance. To determine whether this pattern extended to day-to-day variability *within* mice, we determined CVs across HCM monitoring days for each animal. A similar pattern of variability was observed for all animals and generalized widely across strains (Fig. 3b).

**Figure 3.**
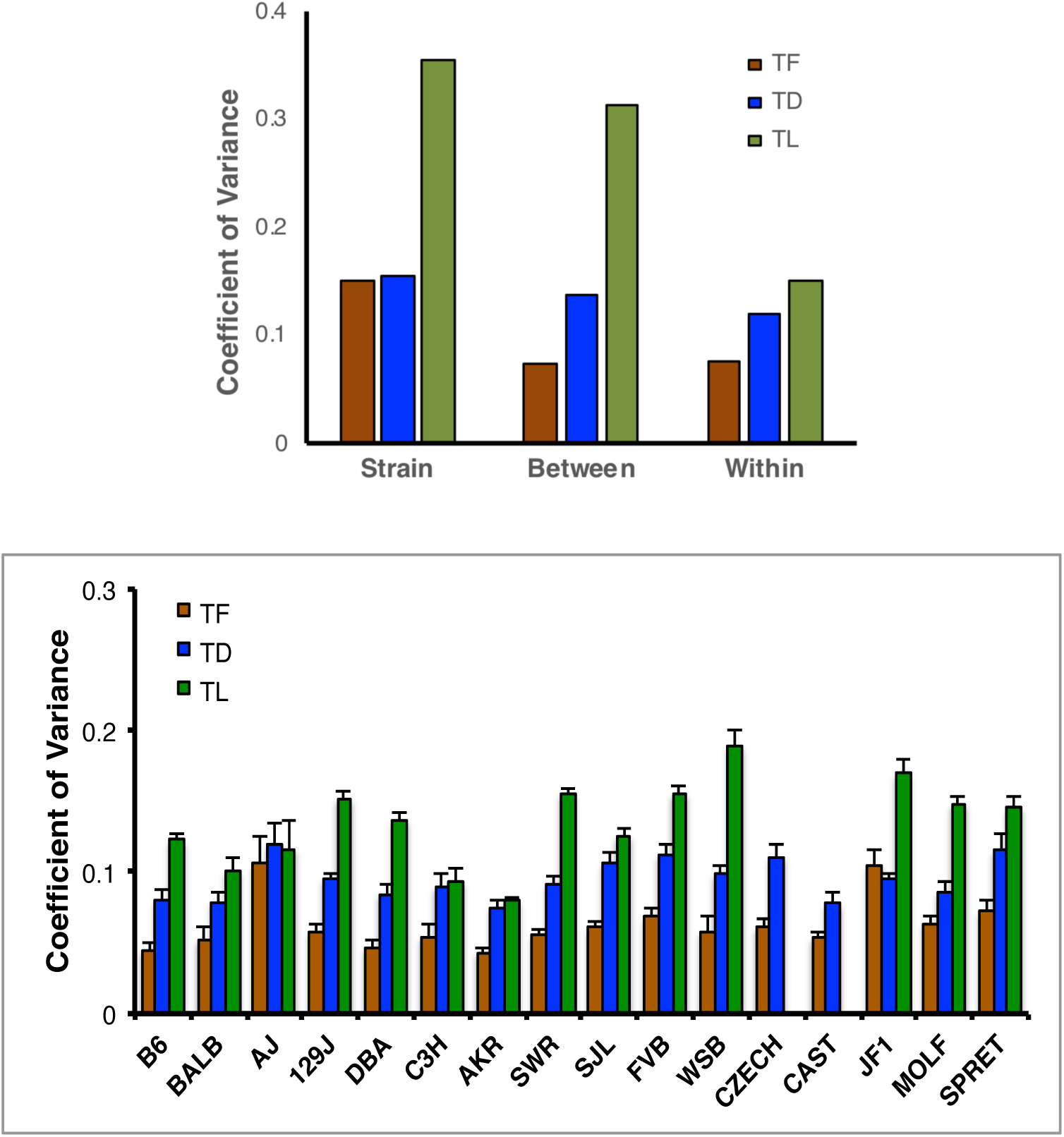
Strain, between-individual, and inter-individual variabilities in food/water intake and locomotion. **(a)** Coefficients of variance are presented for strain averages (Strain), within-strain individual mouse averages (Between), within-mouse day-to-day values across multiple experimental days (Within). Bars are color-coded for food (brown), water (blue) and locomotion (green). **(b)** Coefficients for variance (across experimental days) for within-mouse features plotted strain-by-strain.

### Hierarchically-paired inverse relationships between feeding-related features

To gain insights into how behavioral model features interact to shape patterns of ingestive behavior, we examined correlations among all 24h-averaged feature values from individual mice. This revealed a pattern of significant inverse correlations between behavioral features at each tier of the model for feeding and drinking behavior (Fig. 4). For feeding drinking and locomotor behaviors, relationships between 3 feature pairs (ASI-ASP, BS-ASBR, and BI-BD) are shown in scatterplots, each displaying data from all 169 individual in the study. For feeding and drinking, significant and robust inverse correlations were observed for feature pairs at each hierarchical level. By contrast, analysis of corresponding locomotor features revealed no significant correlations.

**Figure 4.**
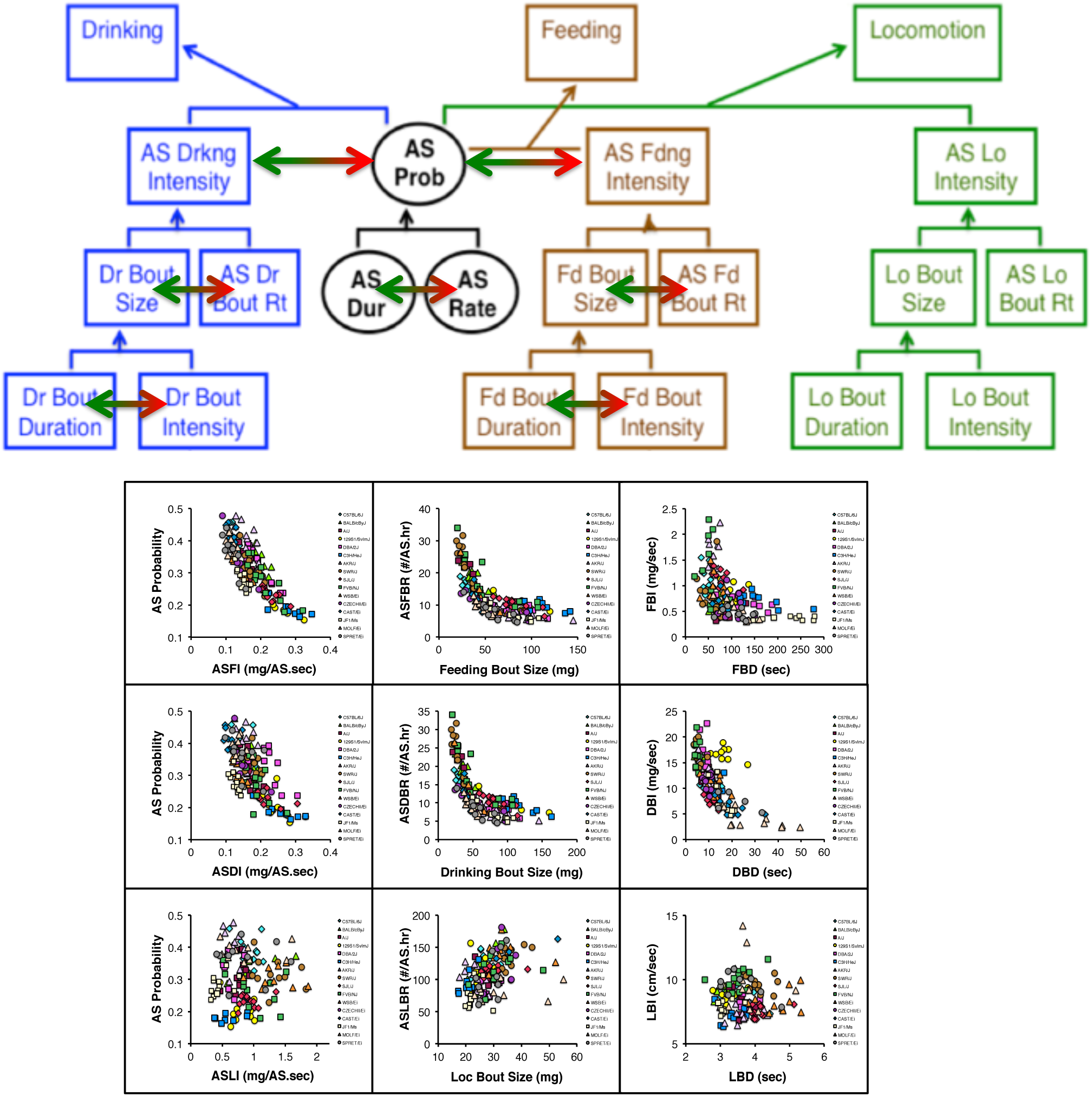
Inversely correlated feature determinants for feeding and drinking. **(a)** At several hierarchical levels, key feature values were determined by inversely correlated features at the level below. Such relationships were observed for feeding and drinking features, but not for locomotor features. **(b)** Scatterplots between corresponding feeding, drinking and locomotor feature pairs are shown below. Robust inverse correlations were observed for ingestive, but not locomotor features.

### Hierarchically-paired inverse relationships between ingestive behavior features are expressed by individual mice

We sought to determine whether the between-mouse relationships seen for feeding-related features were detectable in the longitudinal data generated by individual mice. Fig. 5 displays individual mouse-day data for the C57BL/J, DBA/2J and BALB/cBy strains. In the top row, data are color coded by strain, and below that, data from each of the strains are plotted separately, and color-coded by individual mouse. It is readily apparent that reciprocal correlations between paired feature values at multiple hierarchical levels characterize day-to-day variability within individual mice.

**Figure 5.**
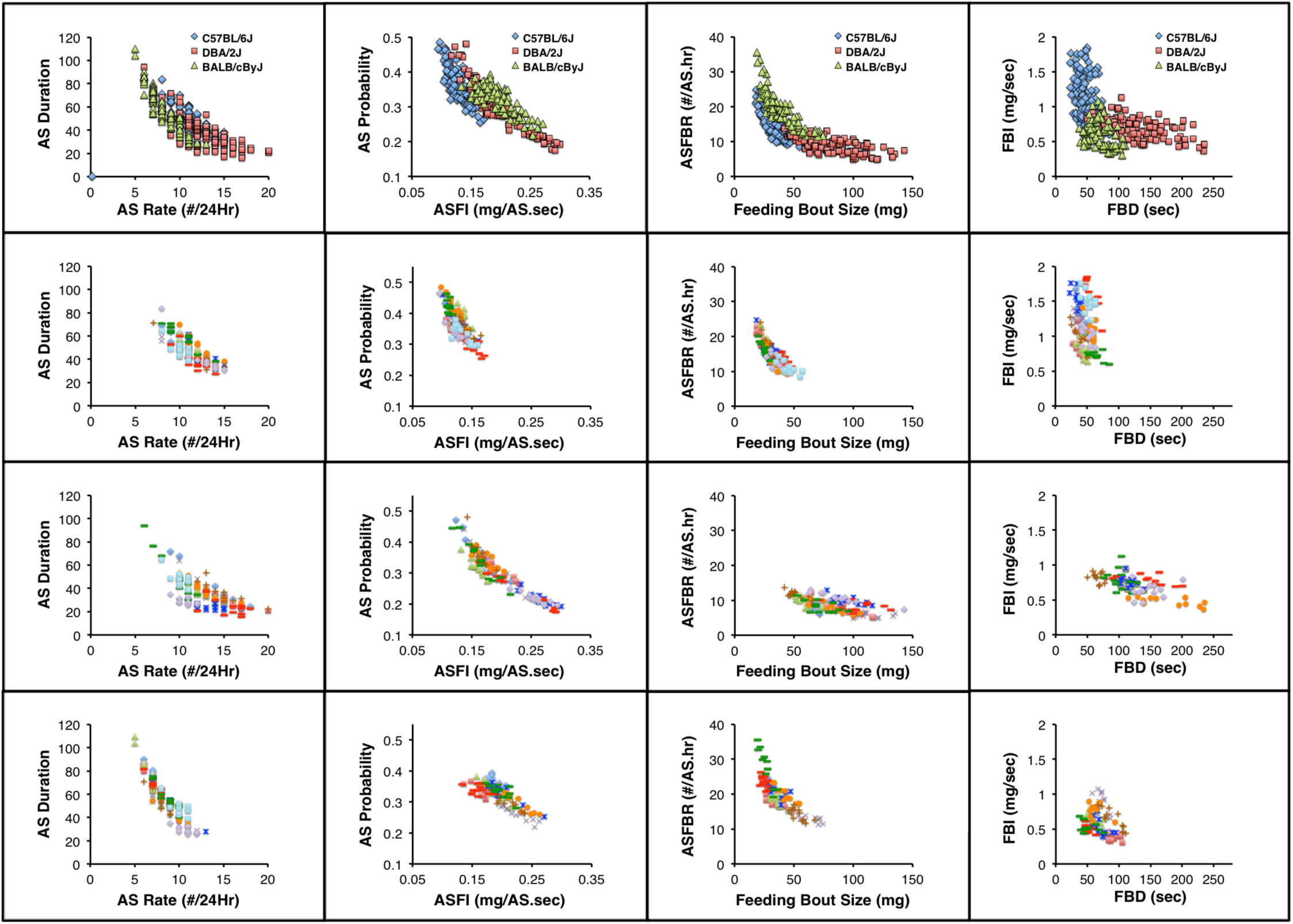
Day-to-day within-mouse variability for Active State, feeding and drinking features exhibited similar inverse correlation patterns. Each data point in this figure corresponds a feature value from a single mouse on a single day. In the top row, scatterplots of feature pairs are presented using all mouse-days for the C57BL/6J, DBA/2J and BALBcByJ mice. The datapoints are color-coded by strain. Each of the 3 rows below display data from one of the 3 strains, color coded by individual mouse.

Although the definitions of locomotor features were arithmetically analogous to those for feeding and drinking, the interrelationships among locomotor feature values differed markedly from those of ingestive behavior. No significant inverse correlations among locomotor features were observed. Instead, patterns of positive correlations were generally observed within and between features at each hierarchical level. These pattern differences between features for locomotion vs. feeding/drinking raise the possibility that the latter may provide a means through which behavioral regulation contributes to the known tight homeostatic regulation of energy and volume balance.

### Ingestive behavior within-animal feature variability decreases at successively higher hierarchical levels

We speculated that a pattern in which particular features are determined by reciprocally related subfeatures may reflect a means for reducing feature variability. We therefore examined within-animal feature variability for feeding, drinking and locomotion features arising from reciprocally related determinants at each of 3 hierarchical levels: Amounts (AS Probability/AS Intensity), AS Intensities (Bout Size/AS Bout Rate), and Bout Sizes (Bout Duration/Bout Intensity) (Fig. 6a). Coefficients of variance for these features across mouse-days were determined for each mouse, and averaged values for all animals in the study are seen in Fig. 6b. In accord with our hypothesis ingestive behavior feature variabilities were lowest for amounts, intermediate for AS Intensities and highest for bout sizes. By contrast, locomotor variabilities were highest for locomotor distance, and least for locomotor bout sizes. These patterns were consistent and robustly generalized among strains Fig. 6c-e.

**Figure 6.**
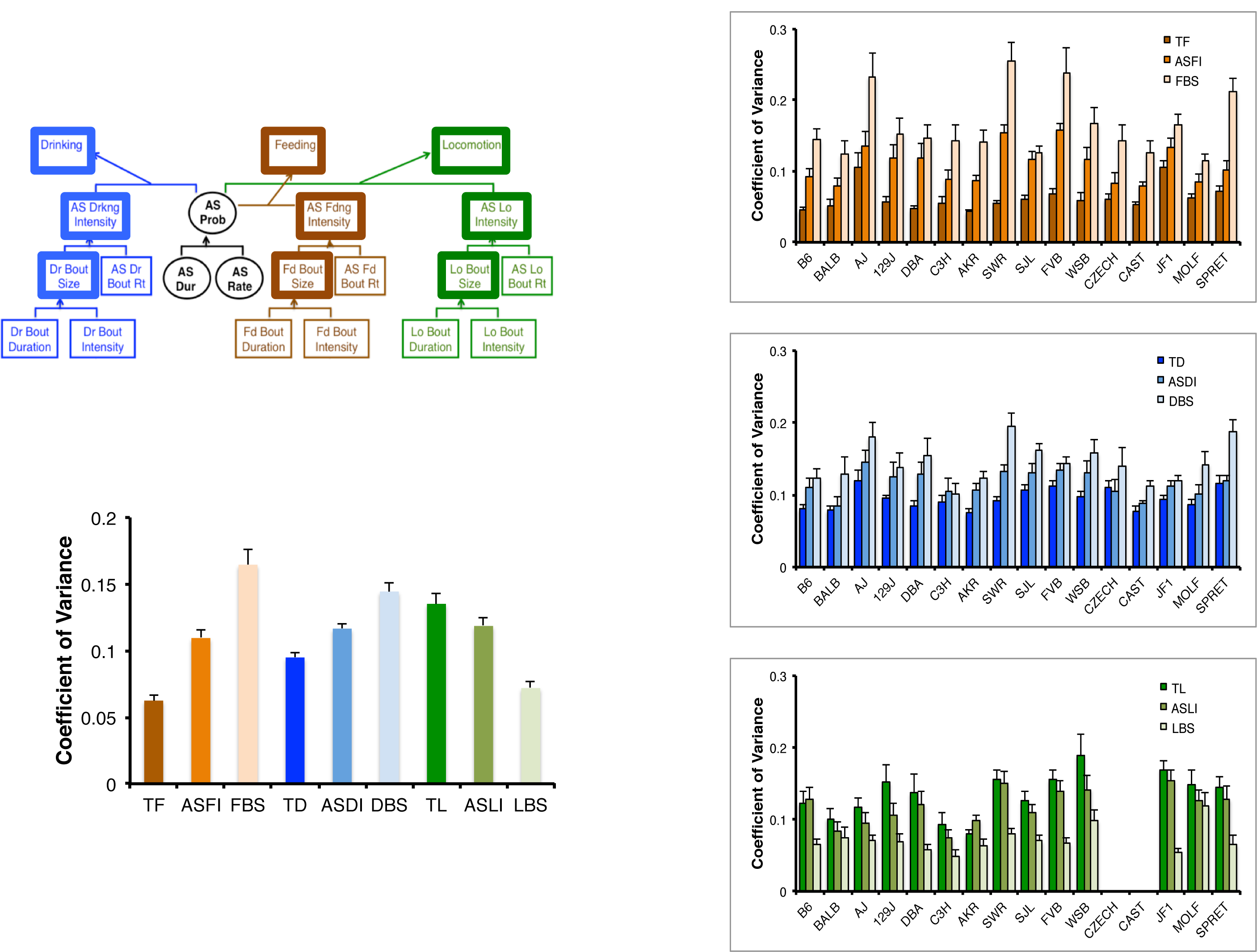
Within-mouse feature variance across hierarchical levels for feeding/drinking and locomotion. **(a)** The features examined were amounts of feeding/drinking and locomotion (top tier), Active State Intensities (second tier), and bout sizes (third tier). **(b)** Coefficients of feature variance for all animals, **(c-e)** Strain-by-strain coefficients of variance for feeding (Fig. 6c), drinking (Fig. 6d) and locomotion (Fig. 6e).

### Buffering daily food intake from low-level behavioral perturbations

We speculated that occurrence of inverse relationships between feature determinants at multiple hierarchical levels would be consistent with a buffering function shielding homeostatically critical output measures (food and water intake) from lower level behavioral perturbations Fig. 7a. Correlation analysis was performed to determine the extent to which gross levels of feeding, drinking and locomotion were sensitive to variation in their corresponding bout properties. For feeding and drinking, gross intake values were significantly correlated with active state feeding and drinking intensities, but that was not the case for features at lower hierarchical levels (Fig. 7b). By contrast, significant and robust correlations were observed between total locomotion and all locomotor bout properties.

**Figure 7.**
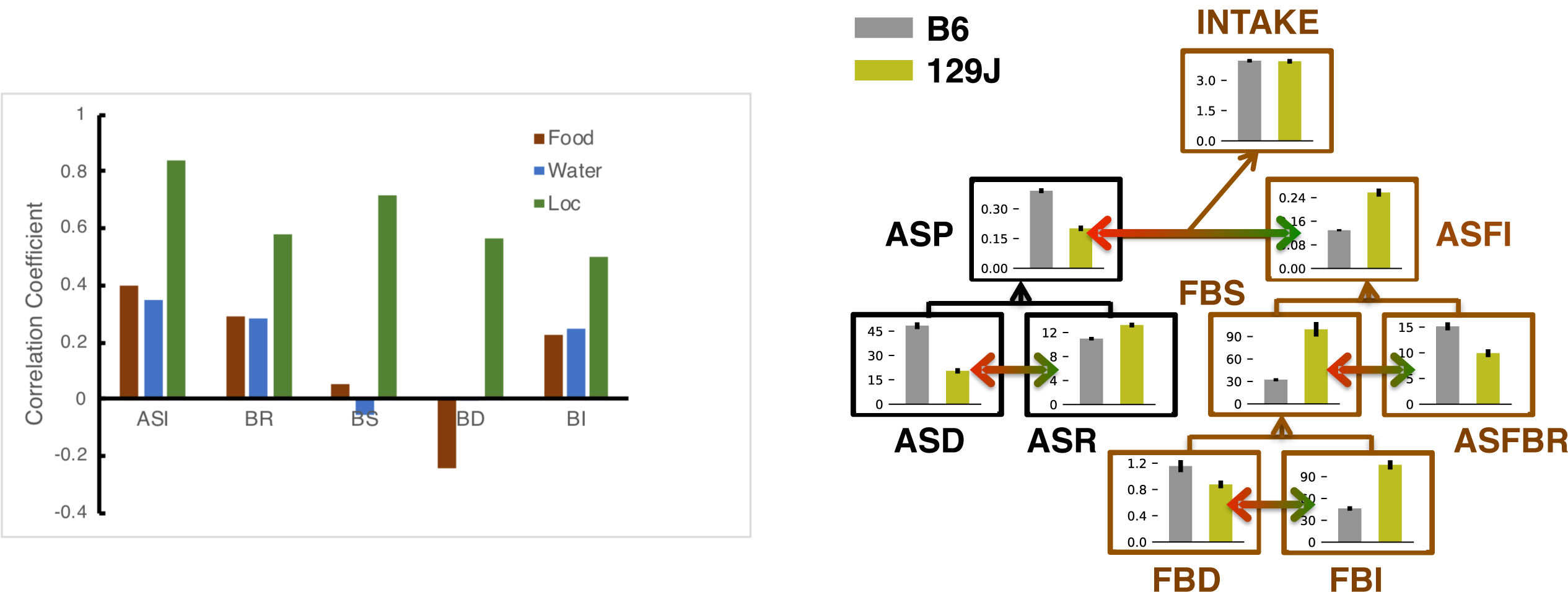
Correlations between top-tier features and bout properties for feeding, drinking and locomotion. **(a)** Pearson correlation coefficients determined between a top tier feature and corresponding Active State Intensities (ASI), bout rates (BR), bout sizes (BS), bout durations (BD) and bout intensities (BI). Significant correlations indicated by asterisks. (b) Active State and bout feature values for 2 strains (C57BL/6J and 129J) with similar total food intake levels despite 3-fold differences in feeding bout size.

A comparison between the C57BL/6J and 129Sv/J strains provides an instructive example of how the above-described patterns of feature relationships may act to blunt the impact of low-level genetic perturbations on gross levels of daily food and water intake. Although these strains exhibit 3-fold differences in their feeding bout sizes, they exhibit nearly identical body weights and levels of food intake (Fig. 7c). However, differences in the resulting AS feeding intensities are partially blunted by reciprocal phenotypic differences in AS feeding bout rates. In turn, the remaining phenotypic differences in AS feeding intensity are then eliminated at the level of gross food intake by reciprocal phenotypic differences in AS probability.

## DISCUSSION

A 2017 WHO report attributes the escalation in the worldwide prevalence of obesity— nearly 3-fold since 1975—to increased intake of foods high in dietary fat and to increasingly sedentary lifestyles^13-15^. This reflects the consensus that in common forms of human obesity, dysregulation of behavioral determinants of energy balance plays the most prominent role in obesity initiation, more so than perturbations that alter metabolic rate or storage. The obesity epidemic results primarily from excessive exposure to nutritional and environmental factors that promote positive energy balance via changes in behavioral control. Understanding the fine-grained patterns of feeding behavior and physical activity are core to effective solutions.

Toward that end, we quantified HCM data for 16 genetically-diverse inbred strains (168 individuals) and applied to it a hierarchically-organized system for classification of endophenotypes relevant to behavioral determinants of energy and volume balance: patterns of feeding, physical activity, and drinking. We discovered a striking pattern of inverse correlations between pairs of ingestive (but not locomotor) bout features whose products determined feature values at the next higher hierarchical tier. The repeated occurrence of such relations at multiple hierarchical levels raises the possibility that they reflect mechanisms for the regulation of behavior in the service homeostatic control.

The centrality of effective energy balance regulation to survival has been a driving force in natural selection, and in mammals, the process is believed to be highly robust: i.e. able to maintain function in a manner resistant to a wide range of environmental and internal perturbations. It has been proposed that this accounts for a frequently-encountered phenomenon in mouse models: genetic perturbations of gene products considered to play major roles in the regulation of feeding behavior often produce no, or modest influences on food intake and body weight^1,16^. This has been suggested to be the result of functional redundancy in neural pathways regulating energy balance. Such a property could enhance species survival: for example, compensatory mechanisms (backup circuits) could enable animals to survive genetic perturbations producing malfunction of a particular energy balance circuit. The patterns of endophenotype expression reported here for ingestive behaviors associated with homeostatic control^17^, and its contrast with patterns associated with locomotion led us to consider several organizational principles common to robust biological systems, including: 1) System control, 2) Redundancy, 3) Decoupling, and 4) Modularity.

*System control* refers to effector mechanisms such as negative feedback that are designed to adapt to a diverse array of inputs by producing a consistent output. For example, the impact of abnormalities in feeding bout sizes are blunted by opposing reductions in feeding bout rates. *Redundancy* refers to the ability of a component of a respond adaptively to the failure of another. For example a dysregulation of bout size and/or frequency could lead to an abnormal AS feeding intensity. The strong reciprocal connections between this measure and AS probability indicate that changes in AS regulation can compensate such bout level perturbations. *Decoupling* refers to the extent to which system output is resistant to variability in low-level features. Our observation that food and water intake levels (by contrast with locomotor activity levels) do not correlate significantly with low-level features is consistent with this feature of robust systems. Finally, *modularity* refers to the repeated occurrence of functionally homologous elements throughout a system. It is possible that this principle is reflected by the repeated occurrence of inversely-related behavioral features at several hierarchical tiers for both feeding and drinking behavior.

Altogether, these findings provide evidence that HCM behavioral dissection not only enhances the ability to detect experimental influences on ingestive behavior, it also reveals how compensatory behavioral processes mobilize to minimize the impact of those effects. These results also illustrate that the most prevalent measure used for studies of feeding behavior—food intake—is a relatively insensitive indicator of experimental influences on feeding behaviors. These findings indicate that a wealth of phenotypic variability in the regulation of ingestive behavior may go undetected when traditional feeding phenotyping methods are used. The “behavioral dissection” made possible by our HCM approach may therefore greatly enhance the extent to which the impact of genetic, pharmacological and environmental (eg: diet) experimental perturbations can shed light on behavioral contributions to homeostatic regulation.

## METHODS

### Animals and Data Collection

Sixteen genetically diverse inbred strains of mice were obtained from the Jackson Laboratory, including strains in common use: C57BL/6J, BALB/cByJ, A/J, 129S1/SvImJ, DBA/2J, C3H/HeJ, AKR/J, SWR/J, SJL/J, FVB/NJ, WSB/Ei, CZECHII/Ei, CAST/Ei, JF1/Ms, MOLF/Ei, and SPRET/Ei. Animals were housed under a standard 24 hour light/dark (LD) cycle, consisting of a 12 h day (150 lux overhead illumination) and a 12 h night. Room temperature was 20°-22°C, and mice had *ad libitum* access to water and standard chow (PicoLab Mouse Diet 20, Purina Mills, Richmond, IN). Animals were acclimated to these vivarium conditions for at least 7 days prior to behavioral monitoring. Male mice approximately 3 months of age were examined, with group sizes ranging from *n* = 9 to *n* = 12 per strain. Experiments were performed in accordance with guidelines of the UCSF Institutional Animal Care and Use Committee.

### Data Collection

Mice were individually housed and monitored for 16 days in HCM cages, as previously described ^12^. As previously described, a four day acclimation period to HCM housing was provided, and the data collected during the subsequent 12 days were used for analysis. Data were collected continuously across days except for a daily maintenance period (Zeitgeber hours 6-8), during which food and water were measured/replaced in a manner that did not require animal handling or opening of cages, minimizing disruption. As previously described, quality control algorithms were run to correct activity platform location drift error, and occasional instances of device malfunction. Data collected during periods of device malfunction were excluded from subsequent analysis. The detection of behavioral events, their organization into ASs and ISs, and the derivation of AS Intensity (ASI) and AS Duration (ASD) features were performed as previously described ^12^.

### Ingestive Bout Designation

Behavioral records consist of time intervals for food and water consumption as well as distance and position coordinates (t, x, y), recorded with a 0.02 second temporal resolution. Feeding behavior is indicated by the temporal pattern of breaks in a photobeam positioned within the HCM feeder assembly at an opening through which animals access powdered food. At the end of each experiment day, the amount of food consumed is manually measured and a feeding coefficient FC [mg/s] is calculated (daily) for each mouse as the total amount of food consumed divided by the total photobeam-breaking time. In an analogous manner, water drinking data is acquired by a lickometer device that measures the onset and offset times of individual lick. A licking coefficient LC [mg/s] is calculated daily by dividing measured 24 hr water intake by the cumulative 24 hr lickometer activation time. Consecutive feeding or drinking events are considered to be contained within the same bout when their event are separated by a time interval smaller than a pre-determined time threshold (Feeding Bout Threshold (FBT) or Drinking Bout Threshold, DBT). The durations of feeding or drinking bouts are designated as time intervals spanning the initiation of the first event until the termination of the last event of each cluster. Optimal hyper-parameters have been found for the Strain Survey Study: FBT = 30s, WBT = 30s, MBVT = 1cm/s, MBDT = 5cm.

### Ingestive bout parameters

Ingestive bout parameters are automatically derived for three sets of time bin lengths: 24 hours, 12 hours (corresponding to the LC and DC) and 2 hours (6 LC bins and 6 DC bins). The parameters consist of:

<u>Active State bout rates</u> for feeding and drinking (ASFBR, ASDBR, respectively) are defined as number of bout onsets divided by time spent in Active State and expressed as # bouts per AS.hour.
<u>Feeding and drinking bout sizes</u> (FBS, DBS, respectively) are determined by dividing the amount of food/water consumed within a time bin by the number of bout onsets expressed during that bin. Bout sizes are expressed in mg.
<u>Feeding and drinking bout durations</u> (FBD, DBD, respectively) are designated as time intervals spanning the initiation of the first event until the termination of the last event of each event cluster. Averaged bout duration values are determined for each time bin be dividing the cumulative time during which a bout type is expressed by the number of bout onsets in that bin. Bout durations are expressed in seconds.
<u>Feeding and drinking bout intensities</u> (FBI, DBI, respectively) are determined for each bout by dividing its size by its duration. Bout intensities are expressed as mg/sec.

### Locomotor bout designation

Locomotor data consists of timestamps t and position coordinates (x, y) acquired by a moving platform, and corrected for position drifting. HCM raw data is used to build spells of activity for feeding, drinking and movement. As a first step, the home base area is identified by mouse position data as a single/double cell in a 2×4 grid cage discretization arrangement. The home base area is detected as the cell with the largest occupancy time on each given observation day (defined as the total time spent in cell divided by the total HCM recording time in a day). In the event that the largest occupancy time is less than 50% of the total recording time, the contiguous cell with the second-largest occupancy time is added to the home base area if the former accounts for at least 25% of the total recording time. Movement occurring in the home base area are not regarded as movement proper, rather as non-locomotor movement, and thus excluded from the movement (locomotor) bouts designation process.

For each mouse displacement between coordinates (x_t, y_t) and (x_\{t+1\}, y_\{t+1\}), the velocity and distance traveled are computed. Movement bouts are identified as those consisting of consecutive displacements with a velocity larger than a pre-defined velocity threshold (Movement Bout Velocity Threshold, MBVT [cm/s]) which result in a distance traveled between bout start-(A) and end-point (B), larger than a pre-defined distance threshold (Movement Bout Distance Threshold, MBDT [cm]). In the event the geodesic (i.e. straight - in a beeline) A-B distance is smaller than MBDT, the sequence of displacements is nonetheless designated as when the total distance traveled, or sum of all displacements, between A and B is still larger than the distance threshold MBDT.

Finally, movement bouts whose endpoints are closer in time than an arbitrarily short interval threshold (Movement Bout Time Threshold, MBTT [s], chosen equal to 0.2s), are connected. These very short interruptions in designated bouts are likely to be related to mouse climbing over walls, resulting in a wide tilting of platform which in turns records large displacements in very short times, and thus high velocities, with the mouse actually remaining in place.

---

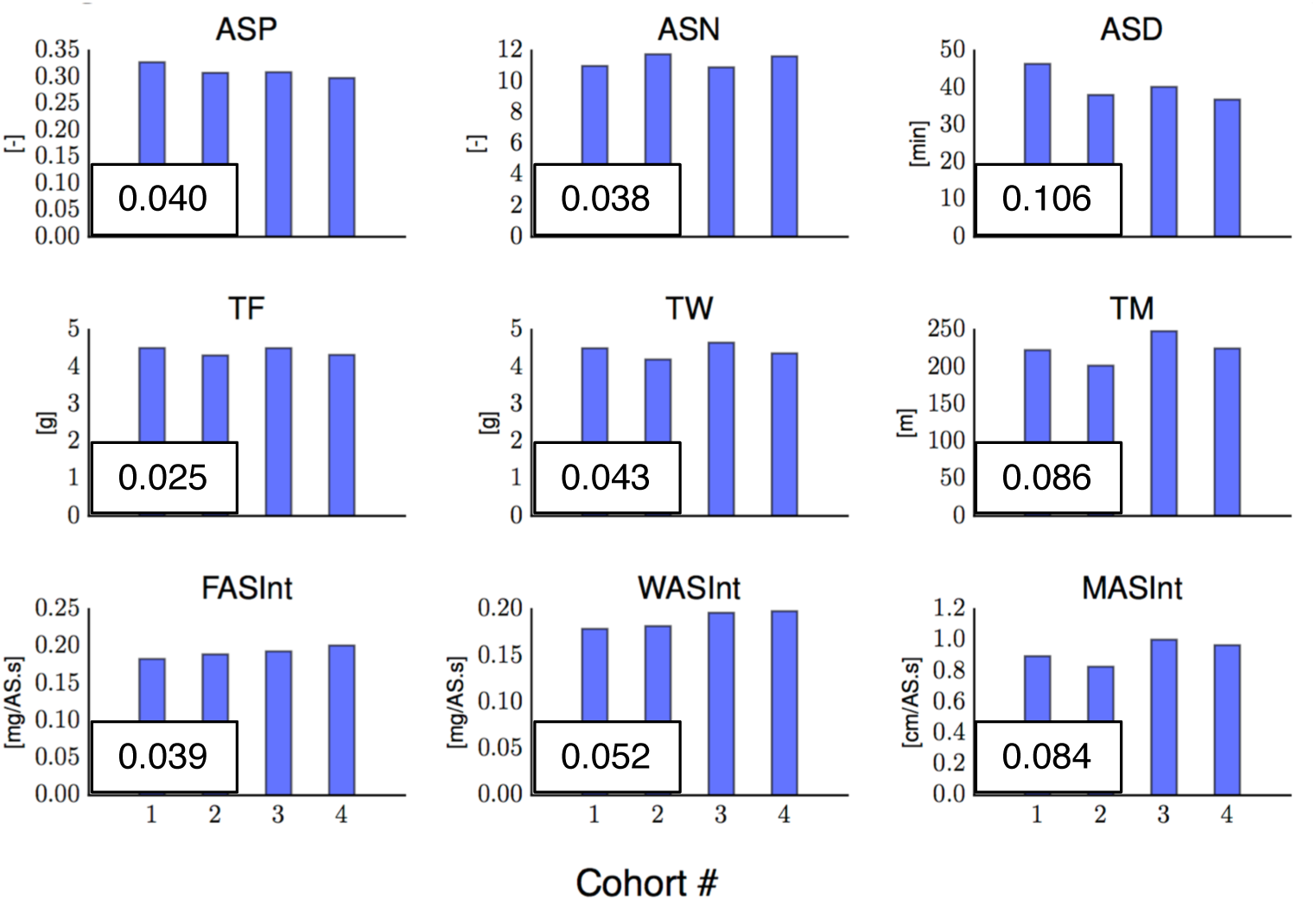

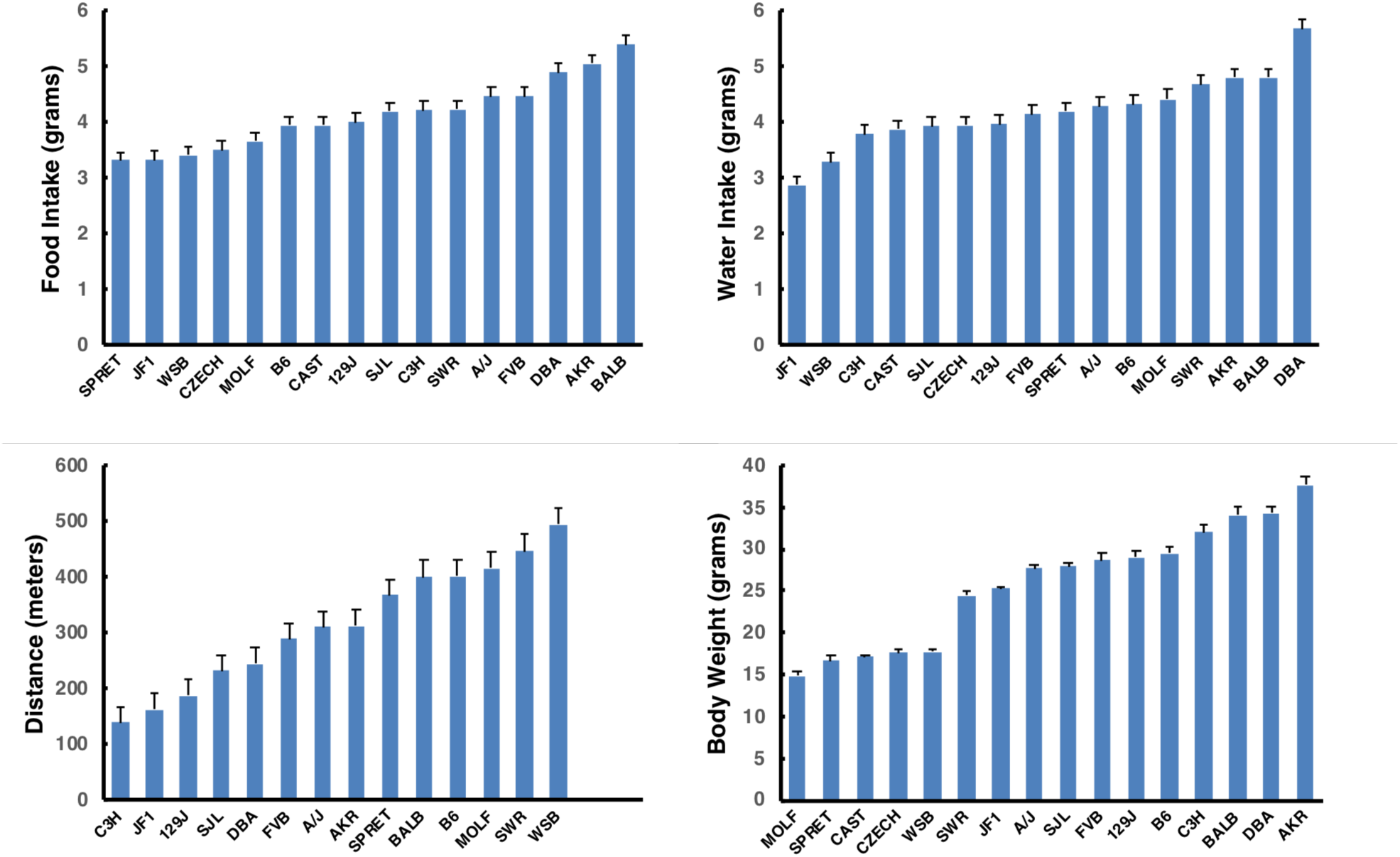

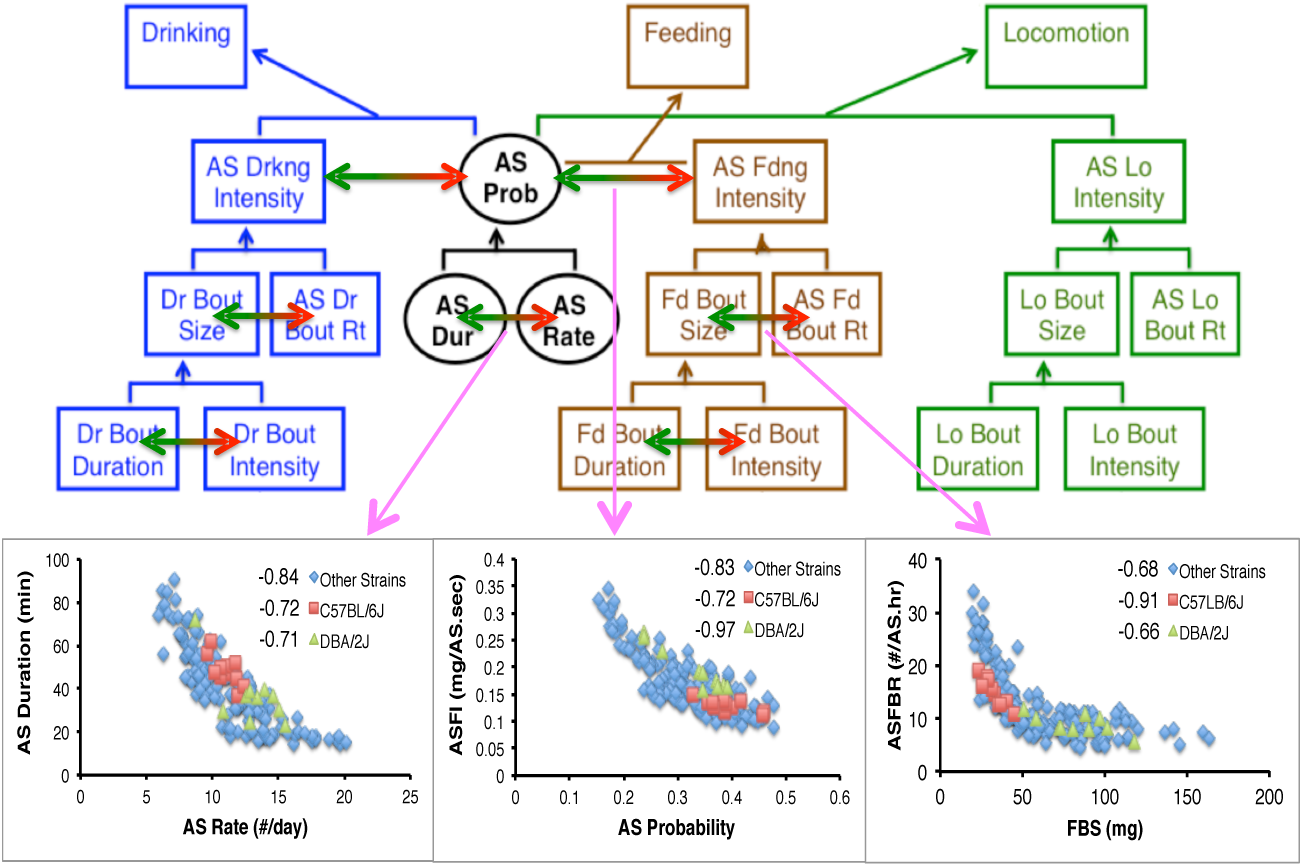

